# Growth increases but regeneration declines in response to warming and drying at Arctic treeline in white spruce (*Picea glauca*)

**DOI:** 10.1101/2023.01.12.523811

**Authors:** Johanna Jensen, Natalie Boelman, Jan Eitel, Lee Vierling, Andrew Maguire, Rose Oelkers, Carlos Silva, Laia Andreu-Hayles, Rosanne D’Arrigo, Kevin L. Griffin

**Affiliations:** Dept of Ecology, Evolution, and Environmental Biology, Columbia University, New York, NY, USA; Lamont-Doherty Earth Observatory, Columbia University, Palisades, NY, USA; Department of Natural Resources and Society, College of Natural Resources, University of Idaho, Moscow, ID, USA; McCall Outdoor Science School (MOSS), College of Natural Resources, University of Idaho, McCall, ID, USA; Department of Earth and Environmental Sciences, Columbia University, Palisades, NY, USA; Tree-Ring Laboratory, Lamont-Doherty Earth Observatory of Columbia University, Palisades, NY 10964, USA; Forest Biometrics, Remote Sensing and Artificial Intelligence Laboratory (Silva Lab), School of Forest, Fisheries, and Geomatics Sciences, University of Florida, PO Box 110410, Gainesville, FL 32611, USA; CREAF, Bellaterra, Barcelona, Spain; ICREA, Pg. Lluís Companys 23, Barcelona, Spain

**Keywords:** Arctic treeline, white spruce (*Picea glauca*), water-limitation, radial tree growth, population regeneration, age structure analysis

## Abstract

As a temperature-delineated boundary, the Arctic treeline is predicted to shift northward in response to warming. However, the evidence for northward movement is mixed, with some sections of the treeline advancing while others remain stationary or even retreat. To identify the drivers of this variation, we need a landscape-level understanding of the interactions occurring between climate, tree growth, and population regeneration. In this study, we assessed regeneration alongside annual tree growth and climate during the 20th century. We used an age-height model combined with tree height from aerial lidar to predict the age structure of 38,652 white spruce trees across 250 ha of Arctic treeline in the central Brooks Range, Alaska, USA. We then used age structure analysis to interpret the trends in regeneration and tree-ring analysis to interpret changes in annual tree growth. The climate became significantly warmer and drier circa 1975, coinciding with divergent responses of regeneration and tree growth. After 1975, regeneration of saplings (trees ≤ 2m tall) decreased compared to previous decades whereas annual growth in mature trees (trees >2m tall) increased by 54% (p<0.0001, Wilcoxon test). Tree-ring width was positively correlated with May-August temperature (p<0.01, Pearson coefficient) during the 20th century. However, after circa 1950, the positive correlation between temperature and growth weakened (i.e., temperature divergence) while the positive correlation with July precipitation strengthened (p<0.01, Pearson coefficient), suggesting that continued drying may limit future growth at this section of Arctic treeline. We conclude that while warmer temperatures appear to benefit annual growth in mature trees, the warmer and drier environmental conditions in spring and summer inhibit regeneration and therefore may be inhibiting the northward advance at this Arctic treeline site. Researchers should consider the interactions between temperature, water availability, and tree age when examining the future of treeline and boreal forest in a changing climate.

## 2. Introduction

Biomes and the ecotones, or transitions, between them are expected to shift as individual species respond to climate change (Körner 2012). The Earth’s largest ecotone, the Arctic treeline, extends more than 13,400 km around the circumarctic region (Callaghan et al. 2002, Ranson et al. 2011), and is believed to be delineated by an isotherm beyond which constraints on physiological processes prohibit the growth and establishment of trees (Sveinbjörnsson et al. 2002, Körner 2012, Paulsen and Körner 2014, Körner et al. 2016). As the Arctic is warming at nearly four times the rate of the global average (Rantanen et al. 2022), the treeline-delineating Arctic isotherm is expected to shift poleward (Körner 2012). If treeline species can keep pace with the shifting isotherm, models suggest that individual trees will increase in size (Körner 2012) and treeline may advance 7 - 20 km poleward per year and displace 11% to 50% of Arctic tundra with woody growth (trees and shrubs) by 2050-2080 (ACIA 2005, Callaghan et al. 2005, Pearson et al. 2013, Zhang et al. 2013). The combination of individual growth and population regeneration of treeline at this scale would have profound local, regional, and global impacts on energy balance (Harding et al. 2002), biodiversity (Skre et al. 2002), ecosystem services (Callaghan et al. 2002), biogeochemical cycling (Zhang et al. 2013), carbon balance, and climate-related feedbacks (Wilmking et al. 2006, Pearson et al. 2013).

However, the evidence for such rapid northward advance of Arctic treeline is mixed (Harsch et al. 2009, Van Bogaert et al. 2011, Mamet and Kershaw 2012, Rees et al. 2020, Maher et al. 2021) suggesting a potential delayed response (Körner 2021) and suggesting that temperature alone does not accurately describe treeline position and behavior (i.e., advance, stationary, retreat) (Rees et al. 2020, Maher et al. 2021). For example, in a recent circumarctic meta-analysis of 151 altitudinal and latitudinal treelines, Rees et al. (2020) found that 52.3% of the sites studied showed advancing, whereas 45.6% exhibited no change, and 2.1% showed retreating treeline behavior. They found that this variation in treeline behavior (i.e., advance, stationary, or retreat) was mainly associated with precipitation rather than temperature, during both the growing and non-growing seasons (Rees et al. 2020). Further, evidence of increased individual tree growth due to warming is mixed; some treeline sites have shown increased growth while others have mixed or no response, a difference largely attributed to changes in water regimes (Briffa et al. 1998, Barber et al. 2000, Wilmking and Juday 2005, D’Arrigo et al. 2008). Therefore, when attempting to understand what drives the response of Arctic treeline to climate change, researchers must increasingly consider the influence of temperature alongside variables related to water availability (Rees et al. 2020), such as precipitation, snow and vapor pressure deficit (VPD).

Temperature and water availability impact treeline behavior both at the individual level through tree growth and at the population level through regeneration. The influences of temperature and water availability on tree growth at treeline can vary with tree size due to several physiological constraints in saplings, including higher respiration rates (Griffin et al. 2021) and limited root structures (Smith et al. 2003) compared to mature trees. Here, we consider early life stages as trees < 2 m tall, hereafter referred to as ‘saplings’; conversely, mature trees are ≥ 2m tall (Tranquillini 1979). Mature tree growth at treeline generally increases with increasing temperature (Kozlowski et al. 2008, Körner 2012). However, the correlation between temperature and radial stem growth (i.e., the growth of annual tree rings) is fading in areas of the Arctic treeline across the globe (Camarero et al. 2021), particularly in the boreal forest (Briffa et al. 1998, D’Arrigo et al. 2008, Andreu-Hayles et al. 2011), while the correlation between radial stem growth and precipitation typically strengthens (D’Arrigo et al. 2008). The causes of this phenomenon, known as ‘temperature divergence,’ are still under debate. Some proposals include recent warming and changes in seasonality, global dimming (i.e., decrease in solar radiation in recent decades), and temperature-induced drought stress which, at treeline, would suggest a switch from temperature-limited growth to water-limited growth (D’Arrigo et al. 2008). Therefore, dissecting the nuances of how temperature and water availability influence growth is necessary to understand the future of this ecotone.

In addition to growth, we must investigate how climate change has influenced population regeneration, as treeline advance cannot occur without population growth. Population regeneration is defined here as the cumulative effect of recruitment and mortality of new trees during early life stages (i.e., several years), hereafter, simply “regeneration.” Regeneration is influenced by temperature and water availability (Holtmeier 2009, Brodersen et al. 2019, Holtmeier and Broll 2019). However, investigating treeline regeneration in response to changing temperature and water availability poses several practical challenges that call for the development of novel approaches. Current *in situ* methods which measure forest recruitment, survival, and individual tree growth via plot-level monitoring or age-structure reconstructions are the gold standard. However, these methods are labor-intensive, restricted to relatively accessible portions of the Arctic treeline, and—in the case of plot-level monitoring—require decades of historical data to capture the current effects of climate change on demographic rates. Further, while Rees et al. (2020) provide the most comprehensive plot-level dataset of circumarctic treeline behavior to date, large portions of Arctic treeline have not been studied as they are difficult to access. Products from satellite remote sensing, such as those derived from Landsat, e.g., Global Forest Cover (Ranson et al. 2011, Hansen et al. 2013), offer access to less well-documented sections of treeline but cannot distinguish between tall shrubs and trees and so greatly overestimate treeline advance (Timoney and Mamet 2020). A more accurate estimate can be obtained via high spatial resolution satellite imagery, but such data do not extend far enough back in time to evaluate the status of treeline advance (Rees et al. 2020). Lastly, the methods to observe treeline must be more easily repeatable to actively monitor changes in treeline behavior in response to climate change.

In this study, we present a novel approach that is uniquely adapted to improve understanding of the impacts of climate change on treeline behavior. Our methodology requires minimal *in situ* labor, approximates historical demographic trends with one year of forest structure data (aerial lidar-derived tree height), and estimates forest regeneration on a scale that better reflects the scale of the Arctic treeline (hundreds of hectares and 10^4^ to 10^5^ trees) than plot-level studies (typically tens of hectares and 10^2^ of trees). We implement this method at an Alaskan Arctic treeline site with the following exploratory goals: (1) to reveal qualitative trends in regeneration of a dominant treeline species (white spruce, *Picea glauca*) during the 20th century and evaluate the trends alongside historical climate; (2) to use tree-ring analyses to observe if and how growth of both saplings and mature trees at our study site responded to climate change. We hypothesize that warmer but drier conditions in recent decades have caused trees growing at Arctic treeline become more water-limited. Therefore, we predict that a study of climate, growth, and regeneration over the 20th century will show that growth and regeneration would decrease if conditions became drier; however, if conditions stayed the same or became wetter, growth and regeneration would increase.

## 3. Methods

### 3.1 Study Area

Field data were collected south of the Brooks Range in Alaska (AK), USA. We established a short (5.5 km) north-south transect at the Arctic treeline along the Dalton Highway (67°59’ 40.92’’ N, 149°45’ 15.84’’W, 727 m a.s.l.) (Figure 1). Along this transect, pockets of boreal forest transition into treeless tundra along the floodplain of the Dietrich River and gently sloping lower elevations before tundra vegetation dominates at only slightly higher elevations.

**Figure 1.**
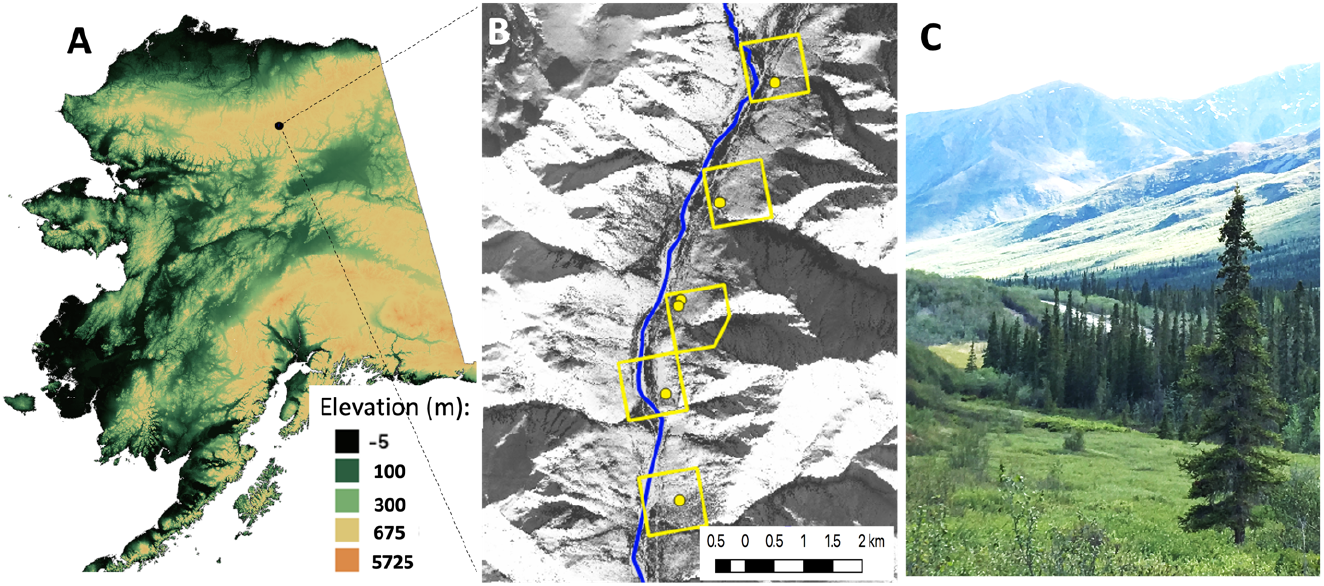
The study area at Arctic Treeline in the central Brooks Range, AK, USA. (A) Elevation map of Alaska with site location. (B) Black and white aerial imagery of the study area in winter, highlighting snow covered tundra (white) and tree cover (black). Yellow dots represent the location of six study plots, the Dietrich River as a blue line and the boundaries of the five forest stands as yellow boxes. (C) Photograph of the northernmost study plot looking south at treeline.

We obtained monthly measurements from 1900-2015 of temperature, vapor pressure deficit (VPD), and precipitation at the study area from the Integrated Ecosystem Model for Alaska and Northwest Canada (McGuire et al. 2016). The average annual temperature from 2000-2015 was −8.25 ± 0.65°C and the average growing season temperature (May – July) was 7.15 ± 0.91°C (mean ± SD). The area is a semi-arid environment (Simpson et al. 2002), receiving 304 ± 77.1 mm of precipitation per year, 45% of which is snow.

Continuous permafrost underlies the Arctic treeline where tree cover predominantly consists of white spruce (*Picea glauca* (Moench) Voss) with occasional black spruce (*Picea mariana* (Mill.) BSP). The understory is dominated by tundra vegetation such as sedges (e.g., *Eriophorum* spp.), moss and lichen (e.g., *Cladonia rangiferina*), short evergreen shrubs (e.g., *Arctostaphylos uva-ursi*), and low (< 1.5 m) deciduous shrubs (e.g., *Salix* spp., *Betula nana* L., *Alnus* spp.). White spruce, our study species, are monoecious and produce cones every year that shed seeds from late August to September (Walker et al. 2012). To the best of our knowledge, there have been no stand clearing disturbances (e.g., fires, insect outbreaks, stand clearing floods) within the study area in the last 200 years, minimizing transient dynamics associated with stand development and succession that otherwise could confound the effects of climate on demographic history (e.g., self-thinning, succession). We found no evidence of bark beetle outbreaks in the field or in tree cores sampled from the area. There are very few standing dead trees (*personal observation*), likely due to the harsh winter winds and thawing permafrost which can topple dead trees.

### 3.2 Summarizing climate conditions during the 20th century

Using the historical climate data, we calculated the monthly precipitation received as snow (SNO) from the monthly temperature and precipitation means; if the mean monthly temperature was ≤ 0°C, the mean monthly precipitation was considered SNO. We use this metric as a proxy for snow water equivalent, although the two are not identical. The annual amount of SNO received was calculated by summing the SNO during the growth year (Aug – Jul). To better understand the overall growing conditions during the 20th century, we used a modified version of the climate index developed by Barber et al. (2000). The climate index differentiates between cool and moist years (positive values) and warm and dry years (negative values) by subtracting the normalized growing season temperature from normalized growth year precipitation. We modified the months included in the growing season to Jun-Aug and growth year to Sep-Aug based on the observed start and end dates of radial stem growth at our sites (Eitel et al. 2020).

### 3.3 Assessing qualitative regeneration at Arctic treeline

We assessed the qualitative regeneration at treeline by first creating a model fitted with *in situ* data that predicts tree age from tree height (Section *‘Age Height Model’*) We then obtained individual tree heights in a 250-ha forest stand using aerial lidar scanning and an individual tree detection algorithm (Section *‘Aerial lidar observations of tree height across forest stands’*). The lidar-derived tree height was then used within the age-height model to predict the age of each tree identified by the tree detection algorithm. Static age structure was constructed by binning the number of individuals in each age class into one-year bins (Section *‘Constructing static age structure’*). We then interpreted the shape of the static age structure to assess the forest regeneration (Section *‘Deducing regeneration from age structure’*)

#### Development of the age-height model from ground observations

Field sampling of tree age and height occurred at six study plots in June 2016 and 2017. All sampled trees were selected within a 20 m radius of a previously established plot center (Eitel et al. 2019). To obtain tree age, a tree core or stem disk was collected from white spruce trees in a random stratified sampling format with five height classes: 0-0.49 m, 0.5-0.99 m, 1-1.49 m, 1.5 – 1.99 m, ≥ 2 m. At each site, one tree from each of the four smallest height classes and three or more trees in the tallest class (≥ 2 m) were selected (n = 70). Additionally, 33 seedlings (≤ 0.3 m) growing in 48 one-meter square quadrants were included (eight quadrants per site, located 10 m from plot center in the cardinal directions; total n = 103).

#### Tree Age

Seedling age was measured by counting the number of whorls (i.e., annual terminal bud scars) on the main stem while in the field (n = 33). To determine the age of mature trees, cores were extracted using increment borers (5.15 mm diameter) and stem disks were collected by felling trees. For trees < 5 cm basal diameter, we collected stem disks at basal height (0 cm) (n = 28). In trees ≥ 5 cm basal diameter, cores were extracted at ~ 25 cm height to accommodate the 16-inch increment borer handle (n = 34). Cores and disks were transported to the Lamont Doherty Earth Observatory’s Tree Ring Lab (LDEO-TRL) for tree-ring measurements. Cores and stem disks were stored in cool, dry conditions before being mounted, sanded with progressively finer sandpaper, and scanned using a high-resolution flatbed scanner (3200 dpi resolution). Tree-ring boundaries were marked digitally on the scanned images to the micrometer resolution (0.001mm) and widths were measured and cross-dated using the computer software programs CooRecorder and CDendro 8.1 (Larsson 2016, Stockton Maxwell and Larsson 2021). The age of a tree was calculated as the number of rings between the innermost ring, or pith, and the ring corresponding to 2016, when height was measured. In cases where the cores did not contain a pith (23% of samples), the number of years to the pith was estimated by using CooRecorder’s digital pith estimator. In these cases, the estimated number of rings to the pith ranged from 1-7 rings. Ring widths were cross-dated against an existing white spruce tree-ring chronology site near Dalton Highway in Alaska (68.5N, −141.63W) (Jacoby and Davi 2000). Cores that could not be cross-dated, either due to very low correlation with the Dalton tree-ring chronology or rot near the pith, were excluded from the study. The final dataset for the age-height model included 94 trees.

We defined ‘tree age’ as a tree’s age in 2016, when height was measured. The ‘year of establishment’ is the year a tree germinated. Because cored trees were sampled at ~25 cm rather than at basal height, the true age of cored trees would have been slightly older than the age at the core’s pith. To compensate for the time it took a cored tree to grow to 25 cm tall, we added 14 years, the equivalent to the mean age of a 25 cm tall tree in an age-height model of only trees sampled at basal height (trees aged via whorls or stem disks; n = 63).

#### Tree Height

Tree height was measured in the field (for trees < 1.5 m tall) or by high-resolution terrestrial lidar scans (TLS) of the study plots taken in June 2016 with a Leica Scan Station C10 (Leica Geosystems Inc., Heerbrugg, Switzerland, see Maguire et al. 2019). The laser instrument has a beam divergence of 0.14 mrad, a scan rate of up to 50,000 points s^-1^, a maximum sample density of < 1 mm, and a maximum range of 134 m at 18% albedo. The nominal distance accuracy is 4 mm and the nominal positional accuracy is 6 mm. Each study plot was scanned from four to six positions to minimize occlusion by vegetation canopies. Three common reflectance targets surrounding each plot were geolocated with a global positioning system (GPS - Trimble R7, Trimble, Dayton, Ohio) and an external Zephyr Geodetic antenna. Both the vertical and horizontal accuracy of the GPS are ± 5 mm (Trimble, Dayton, Ohio). For each plot, scans were georegistered using these common reflectance targets using Cyclone 9.1 software (Leica Geosystems Inc., Heerbrugg, Switzerland). UTM coordinates for each tree were manually extracted from the georegistered TLS point cloud for the top of each tree crown. Crown height for each tree was derived from the georegistered TLS point cloud by manually identifying the top of crown and base of bole laser returns, respectively, using Cyclone 9.1 and calculating the difference of the z-values of UTM coordinates (1 cm resolution).

#### Age-Height Model

Age and height were log-transformed to achieve normality and homoscedasticity prior to fitting a simple linear model. In linear space, the linear model becomes a power function with the form

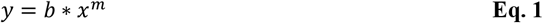

where *y* is tree age, *x* is tree height, and *b* and *m* are model coefficients describing the y-intercept and rate of increase, respectively. To obtain posterior distributions of coefficients *b* and *m*, we fit the linear model to a random subset (75%) of the log-transformed data for 100,000 iterations in R statistical software (R Core Team 2021). Model iterations which were identified as heteroscedastic using a Breusch-Pagan test were excluded from the study (0.4%), resulting in 99,638 final iterations. Model fitting parameters (i.e., r^2^ and root mean squared error, RMSE) were calculated for each model iteration.

#### Aerial lidar observations of tree height across forest stands

An individual tree detection (ITD) algorithm (Silva et al. 2016)identified the location and height of each tree within five ALS tiles containing the six study plots taken by the Alaska Department of Transportation (Hubbard et al. 2011) in May and July 2011 (Figure 1b) covering 250 ha of Arctic treeline. ALS data were collected by a Leica ALS60 sensor mounted on a Cessna Caravan 208Bs aircraft (Hubbard et al. 2011). ALS data were processed using LAStools (Isenburg 2017) in three steps using the lidar point cloud data to ultimately obtain the canopy height model. First, ground returns were classified using progressive triangulated irregular network densification algorithm (Axelsson 2000), to create a one-meter resolution digital terrain model. Second, height above ground was normalized by subtracting the elevation of the digital terrain model. Lastly, a 0.5-m resolution canopy height model was created. ALS-derived 0.5 m canopy height models were smoothed by a 3 × 3 mean filter to remove spurious local maxima caused by tree branches, and individual trees were detected using fixed treetop window size of 3×3 (Silva et al. 2016). In addition, we limited the study to trees ≥ 30 cm tall as is typical in lidar studies elsewhere in at the Arctic treeline (Streutker and Glenn 2006, Naesset and Nelson 2007).

To ensure the trees included in the study were white spruce, as opposed to tall Salix spp., the tiles were visually classified into six land cover types using high resolution (1.8 x 1.8 m) World-View 2 (WV-2) imagery in QGIS (QGIS Development Team 2022): white-spruce-dominated canopy, tundra, rivers, ravines, roads, and pipeline. Only trees within the spruce-dominated canopy were considered in this analysis. To reduce the influence of an anthropologically created edge and/or hydrology on growth rate, trees were excluded from the study if growing within a 20 m buffer of roads (Dalton highway and pull outs) and the Dietrich River (flood plains) as well as those within 15 m of ravines and the Dalton Highway pipeline. Such supervised classification based on collocated imagery of shrubs is commonly done with great success (Timoney and Mamet 2020).

#### Constructing static age structure

To obtain the static age structure of our study area, we used the age-height model (Section *‘Age-Height Model’*) to predict the age of each tree detected in the ALS tiles (Section ‘ *Aerial lidar observations of tree height across forest stands*’). To ensure that the error of the age-height model would not alter our conclusions of regeneration, we propagated the model error into the age structure in the following manner. First, we randomly sampled the posterior distributions of the parameters once (posterior distributions were obtained by fitting the model 100,000 times) and predicted the age of each tree detected in the ALS tile. Then, we constructed age structure by binning the number of trees in each one-year age class. We repeated this process 1,000 times starting with another random sample (with replacement) from the posterior distributions each time to create 1,000 possible versions of age structure at our study area. Then, we averaged the number of individuals in each one-year bin across the 1,000 age structures to obtain a mean and 95% confidence interval of the number of individuals in each age class. Lastly, because aerial tree detection of trees was limited to trees ≥ 30 cm tall (Section *‘Aerial lidar observations of tree height across forest stands*’), the age structure was truncated to the corresponding age (i.e., 16 years according to the age-height model).

#### Deducing regeneration from age structure

We deduced regeneration by observing the shape of static age structure, or the number of trees in each age class at a given time. The age structure in mature, undisturbed forests has many saplings and exponentially fewer old trees (e.g., negative exponential shape). This shape, called a Type III survival curve, is representative of a forest that is continuously regenerating and therefore maintaining or growing the population (Vlam et al. 2017). However, if a regeneration has recently failed to recruit new individuals into the population, the static age structure is unimodal: the drop in older trees is due to accumulated mortality, while the drop in younger trees is due to decreased regeneration relative to previous rates. In this way, the shape of a forests’ static age structure is indicative of the relative rates of regeneration influenced by favorable or unfavorable conditions (Vlam et al. 2017).

### 3.4 Radial stem growth during the 20th century

#### Growth of saplings and mature trees before and after 1975

Circa 1975, climate factors, tree ring chronologies, and tree age structures all underwent notable changes (see Results). Therefore, we further analyzed changes in growth in two periods: before and after the year 1975. To evaluate the degree to which the growth rate of saplings and mature trees varied with tree age before and after 1975, we repurposed the tree cores and disks that were collected and measured for tree age (see Section ‘*Tree Age*’). Growth rate was calculated from the raw ring width as the basal area increment per year, or BAI (cm^2^ yr^-1^). BAI was derived by first calculating basal area over the tree’s lifetime via the area of a circle where the radius is the cumulative sum of ring widths from the pith to the respective year. BAI is then the difference in basal area between the current and previous year. We then developed two generalized linear models with the form

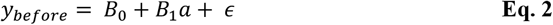

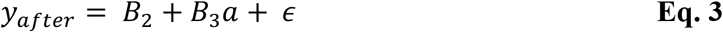

where *y* is growth rate and the subscript indicates *before* or *after* 1975; *a* is tree age; *B_0_* and B2 are the y-intercepts; and *B_1_* and *B_3_* are coefficients.

We assembled separate datasets for saplings and mature trees, and applied both models to each. Both datasets included cores with complete ring records from pith to 2016. Missing rings, which frequently occur in conifers during particularly stressful years (e.g., defoliation or dry growing conditions; see Wilmking et al. 2012a, Leland et al. 2016, Novak et al. 2016), were identified via cross-dating (see Section *‘Tree Age’*) and interpreted as zero growth. We considered a sapling to be < 2 m tall, which is generally equivalent to < 50 years old (see Results, Figure 4a). However, because the oldest trees established after 1975 were 40 years (i.e., oldest in 2016), we limited our analysis of saplings to those ≤ 40 years old. Additionally, we restricted our data set for saplings to basal area grown during the period in which the tree was established to avoid any unintended bias (i.e., we excluded cases where trees established before 1975 grew after 1975). Outliers beyond the 5^th^ and 95^th^ percentiles were removed to reduce noise in evaluating relationships. The final data set for saplings included 59 trees with 1,483 observations of growth rate. For the analysis of mature trees, we selected trees ≥ 50 years old (≥ 2 m tall; see Results) which grew both before and after 1975, resulting in 14 trees with 910 observations of growth rate (age range: 68 to 153 years).

Because growth rate was non-normally distributed, we employed generalized linear models fit by maximum likelihood estimation (Laplace Approximation) using the ‘lme4’ package in R (Bates et al. 2015). We selected the best distribution and link function by comparing the Akaike Information Criterion (AIC) (Akaike 1973) of models with two types of distributions (i.e., Gamma and Poisson) and three types of link functions (i.e., ‘log’, ‘inverse’ and ‘identity’ link functions). For both saplings and mature trees, we selected the gamma distribution with ‘identity’ link function because it had the lowest AIC, and therefore best fit. We added tree ID as a random effect to address the repeated sampling structure of our data (Supplementary Material Table 1). However, since this made the sapling models singular, the random effect was retained only in the mature tree model. We then compared the model equations (Eq. 2 and Eq. 3) within each group (saplings and mature trees). For the sapling models, we used a t-test while the random intercept of the mature tree models required a z-test (Bates et al. 2015). A significant difference between the two models would indicate that the relationship between age and growth rate changed after 1975.

**Table 1.**
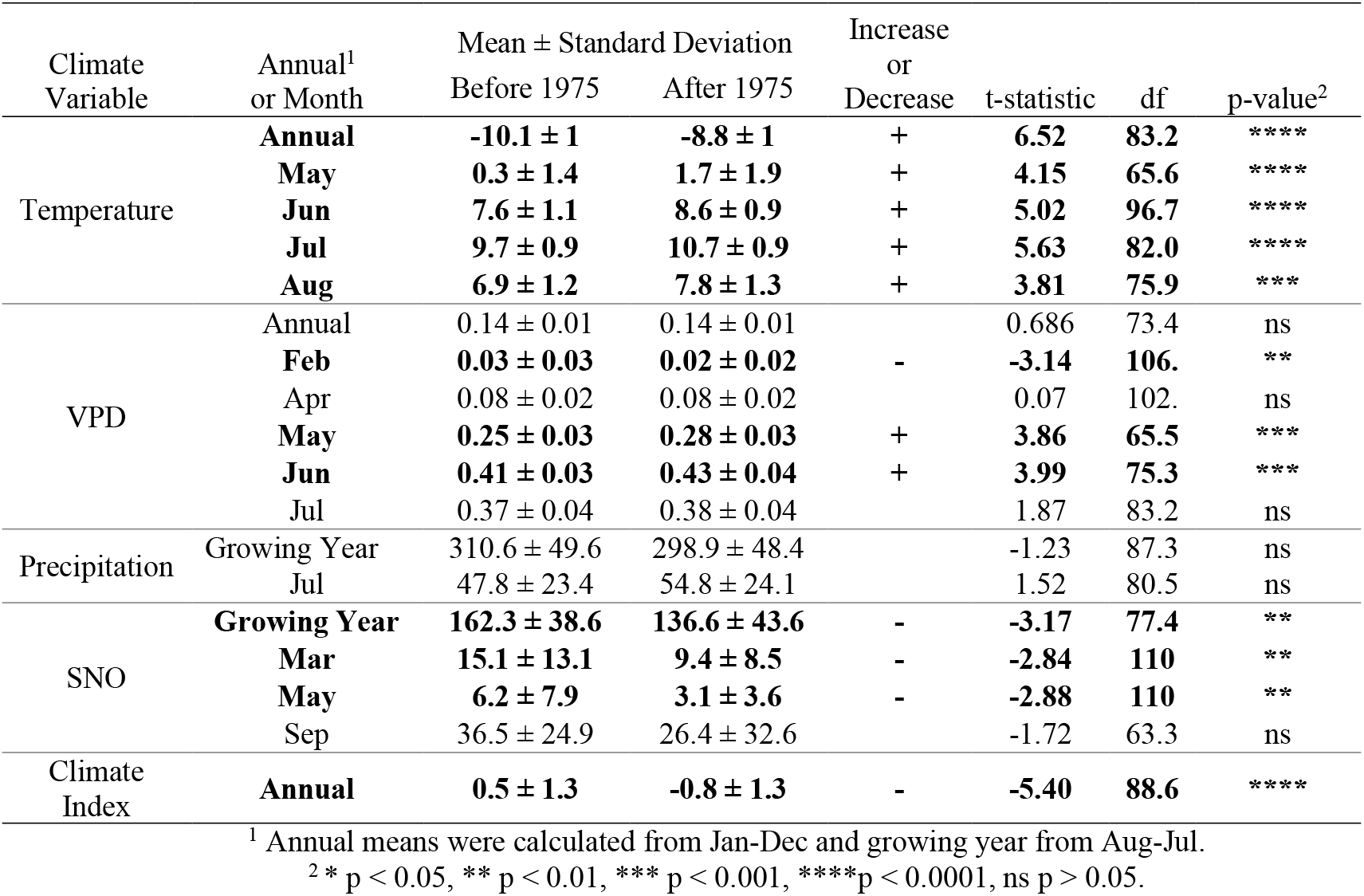
Comparison of annual and monthly climate means at Arctic treeline in central Brooks Range, AK before and after 1975. The selected months are presented because they are significantly correlated with the MNE chronology (TRC; see Table 3). Terms in bold are statistically different after 1975 (p < 0.01, t-test).

#### Relationships between radial stem growth and 20th century climate

We compared the tree-ring width record with meteorological data to elucidate the effects of temperature, precipitation, VPD, and SNO on radial stem growth during the 20th century. To ensure that the trees selected for tree-ring analysis exhibited little-to-no age-related growth trends, we limited our sample to trees whose cores spanned at least 100 years from 1900-2015 (sampled in 2016). The radial-growth sample includes 22 cores from our six study plots, 17 of which cover the entire study period and the remainder covering at least 103 years (mean age 141 years; range: 103 to 232 years). Therefore, this tree-ring analysis is representative of only mature white spruce trees at the Arctic treeline study site.

Raw tree-ring widths were used to build ring width chronologies and then compared to monthly climate during the 20th century. Following standard dendrochronology practices (Cook and Kairiukstis 1990), the ring widths were detrended using two methods: (1) a modified negative exponential (MNE) which removes age-related changes in ring width and (2) a two-thirds spline which ensures a minimum loss of low-frequency variance (Cook 1985, Cook et al. 1995). Using the resulting detrended data, we built two mean tree-ring chronologies (TRC) which provide annual tree-ring width. The detrended data were whitened prior to averaging using an autoregressive time series model. The chronologies were truncated to the year in which the sample depth (number of cores in the chronology) was substantial enough to create a minimum subsample signal strength of 0.85 (Cook and Kairiukstis 1990).

Pearson correlations coefficients, *r*, were calculated from the TRC and monthly climatic data from previous January to current September. Stationary bootstrapping was used to calculate the significance and confidence intervals of each correlation according to Politis and White (2004). Correlations were considered significant at α = 0.01. To elucidate how the climate variables which were significantly correlated with TRC changed over the 20th century, we compared the mean monthly climate before and after 1975 using a t-test.

We used the two chronologies to satisfy different purposes: First, we interpreted the MNE chronology as analogous to that of annual production or growth since the effect of detrending with the MNE method is generally modest in mature trees (Cook and Kairiukstis 1990). This was corroborated by the fact that there was no major difference between the correlations of the MNE chronology and climate and those of raw ring width and climate. Therefore, correlations between climate and the MNE chronology can be interpreted as the correlations between climate and annual growth. The second chronology, a spline, was used to evaluate the stability between TRC, radial stem growth, and climate during the 20th century by computing 31-year rolling correlations between the standard TRC and the mean monthly temperature and precipitation to assess divergence. Tree-ring analyses were facilitated using the R package ‘dplR’ (Bunn et al. 2010) for detrending and building chronologies, and the R package ‘treeclim’ (Zang and Biondi 2015) for conducting climate analyses.

## 4. Results

### 4.1 Climate during the 20th century: Before and after 1975

The climate index revealed a significantly warmer and drier climate after 1975 compared to previous decades (p <0.001; Table 1, Figure 2). For example, from 1900 to 1975, only 37% of the years were classified as warm and dry while the remainder were typically cool and wet. However, from 1975 to 2015, the number of years classified as warm and dry was much higher at 74%. This shift to a warmer and drier climate is reflected in the annual temperature, precipitation, and SNO during the 20th century (Figure 3). In the period after 1975, annual temperature significantly increased by 1°C relative to the period before 1975 (t-test, p < 0.001) (Table 1; Figure 3a). However, growing year (August-July) precipitation remained relatively stable throughout the 20th century (p > 0.05) (Table 1, Figure 3c). The growing year SNO decreased significantly by an average of 25.7 mm after 1975 (p < 0.01) (Table 1, Figure 3d). The drying trend was not reflected in annual VPD (p > 0.05) (Table 1, Figure 3b); however, May and June were significantly drier after 1975 relative to before (Table 1, Figure 3*f*).

**Figure 2.**
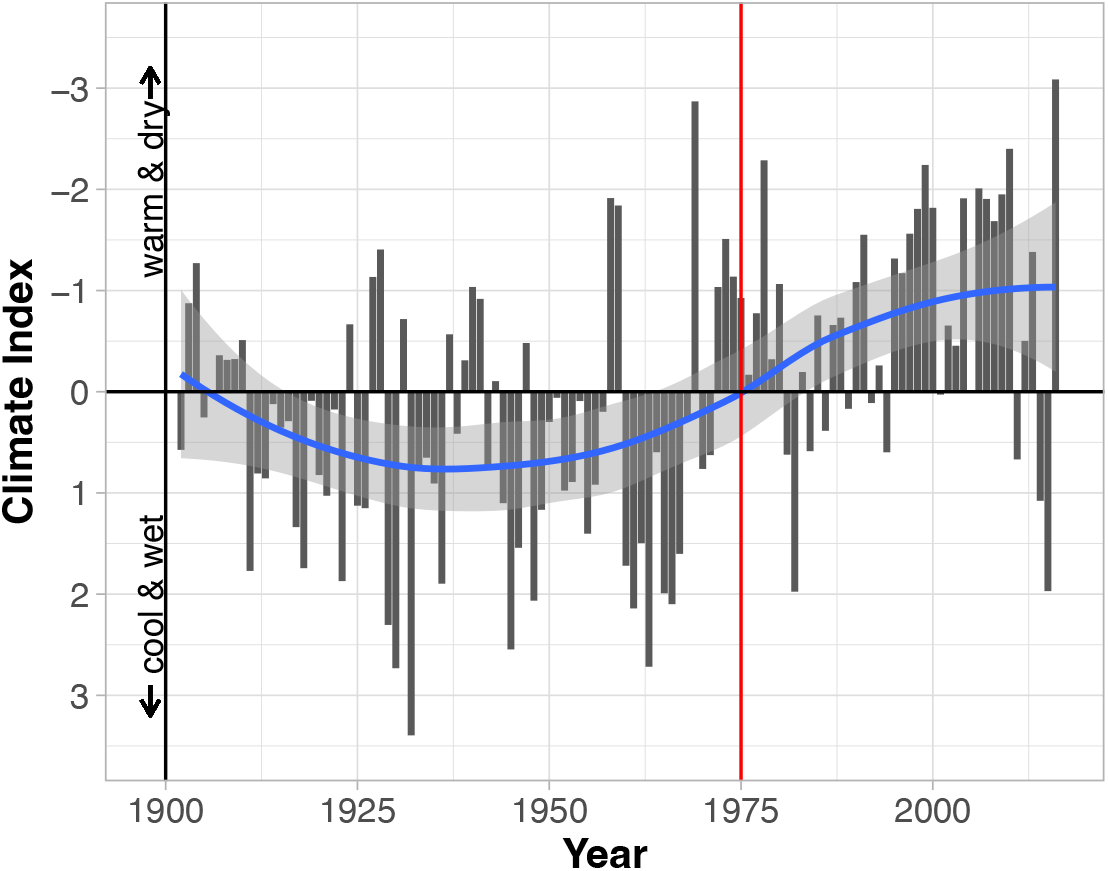
Climate trend from 1900-2016 at Arctic treeline in central Brooks Range using the climate index developed by Barber et al (2000). The index differentiates between relatively cool, wet years (positive values) and warm, dry years (negative values). A loess smoothing curve (blue line) with 95% confidence intervals (grey shading) shows the overall trend through time. The vertical red line indicates the year 1975.

**Figure 3.**
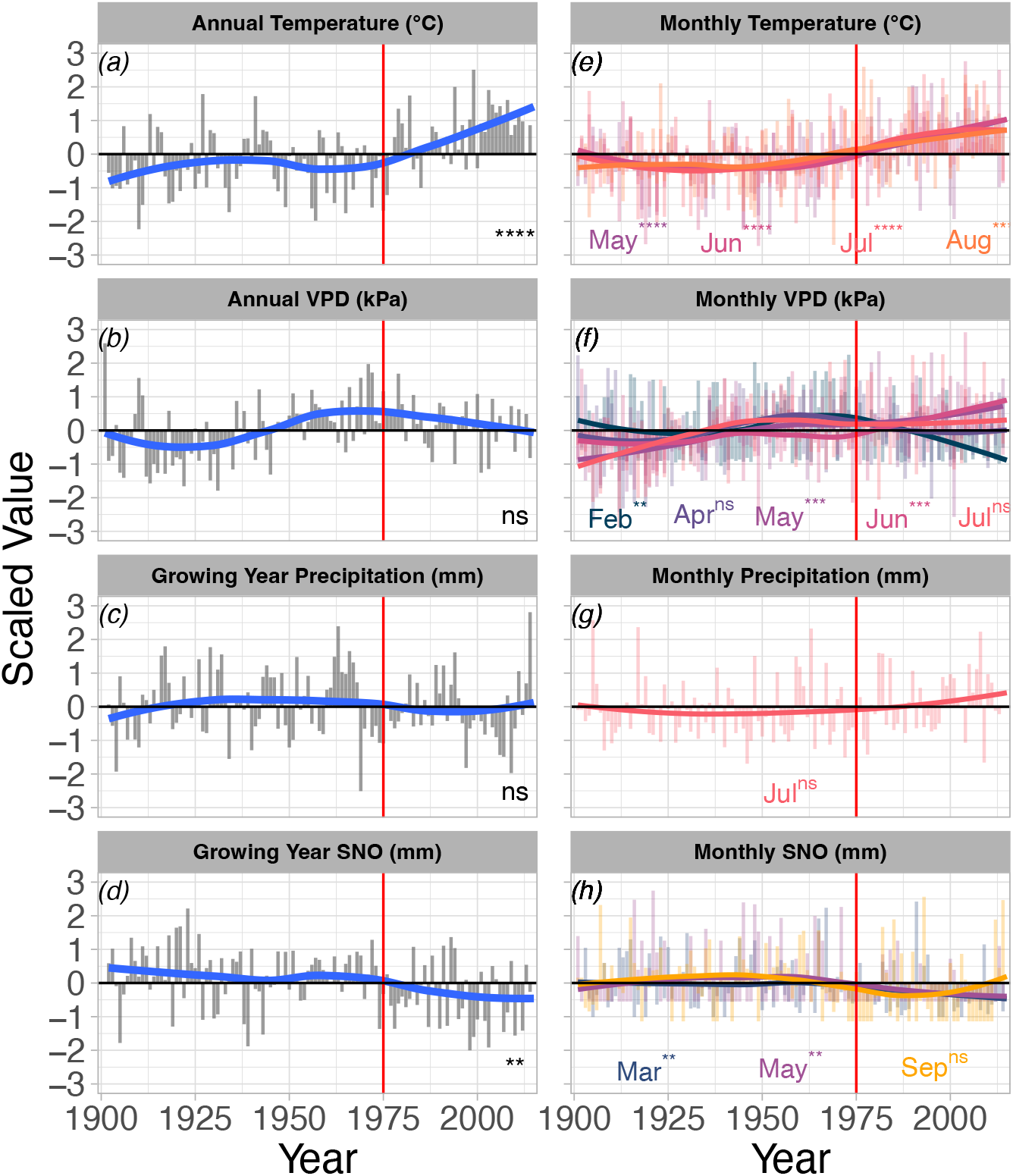
Trends in yearly (a, b, c, d) and monthly (e, f, g, h) climate means from 1900 – 2015 at Arctic treeline in central Brooks Range, AK. Annual means were calculated from Jan-Dec and growing year from Aug-Jul. The selected monthly climate variables are presented because of their significant correlations with tree ring width during the 20th century (see Table 1). Asterisks show p-values from t-tests of the mean before and after 1975 (* p < 0.05, ** p < 0.01, *** p < 0.001, ****p < 0.0001, ^ns^ p > 0.05).The y-axis of all variables is expressed in scaled units where zero is record mean (1900 – 2015) and units are standard deviation. Loess smoothing curves (blue line for annual means; colors for monthly means, legend on each panel) highlight the overall trend through time. The vertical red line indicates the year 1975.

Trends in monthly climate variables are generally similar to annual trends, with the exception of VPD. Mean temperatures during spring and summer (i.e., May, June, July, and August) increased by 0.9 – 1.4°C after 1975 (p < 0.0001) (Table 1, Figure 3e). Additionally, there was no significant difference in July precipitation between the periods prior to and after 1975 (p > 0.05) (Table 1, Figure 3g). SNO in March and May decreased by 38% and 50% after 1975, respectively (p < 0.01), while September SNO remained stable throughout the study period (p > 0.05) (Table 1, Figure 3h). VPD increased in May and June by 12% and 5%, respectively (p <0.001), and decreased by 33% in February (p < 0.01) (Table 1, Figure 3f).

### 4.2 Assessing regeneration: tree age-height relationship and stand-level height and age structure

Figure 4a shows the relationship between age and height in log and linear space. The value of the y-intercept, *b*, and the exponential rate of increase, *m*, were 32.44 ± 0.71 and 0.58 ± 0.01, respectively (mean ± SD). The average r^2^ value of the model was 0.8932 ± 9.90×10^-8^ (mean ± SEM) and ranged from 0.8453 to 0.9293. The RMSE was 23.8 years.

**Figure 4.**
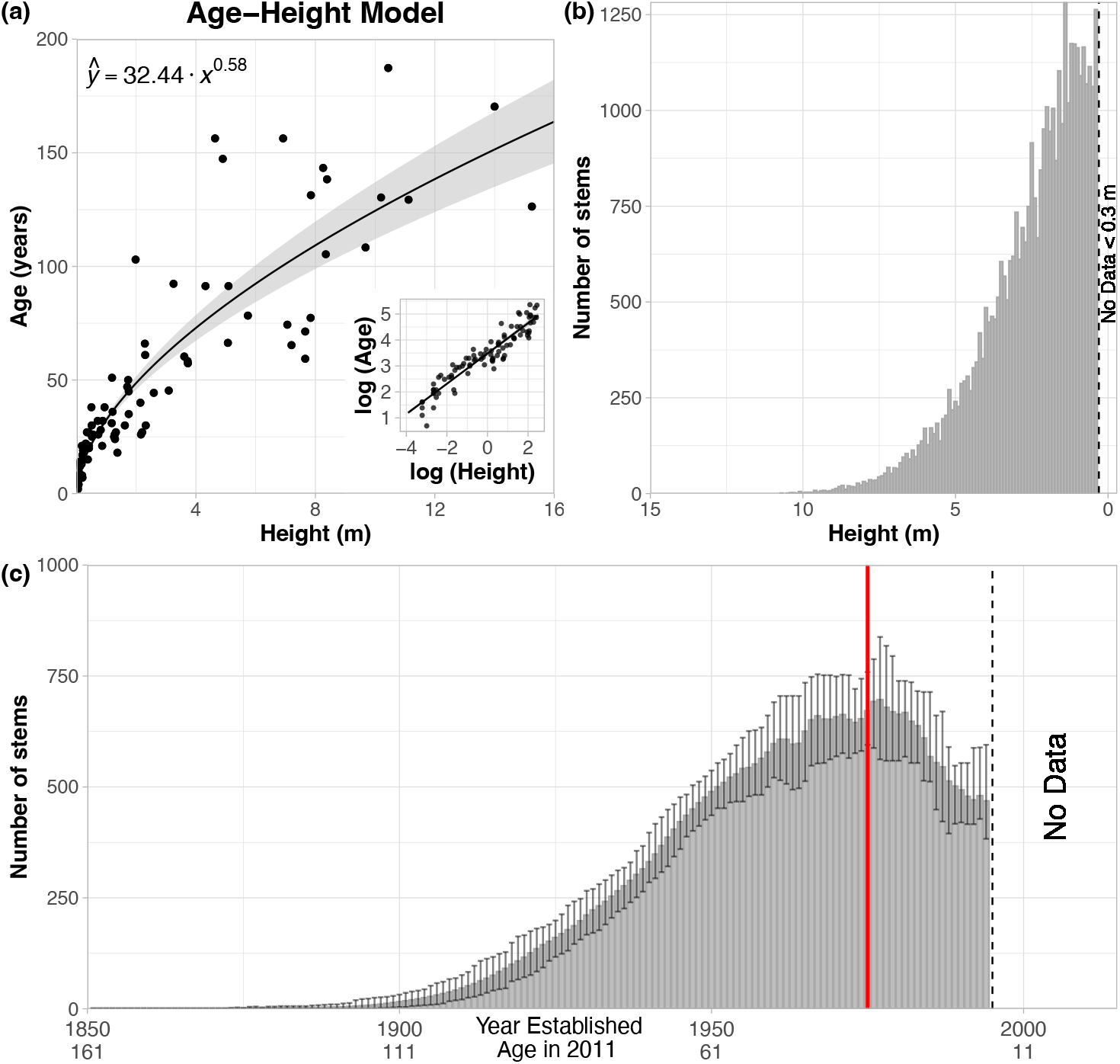
Forest structure at Arctic treeline in central Brooks Range, AK. (a) Relationship between age and height for white spruce at the Alaskan treeline (black line is the mean and grey shading is the 95^th^ percent confidence interval; inset is the relationship in log-log space) (n = 94; mean r^2^ = 0.89). (b) The lidar-derived height structure for white spruce at the study area (~250 ha). Bin widths are each 0.1 m. (c) Age structure of white spruce trees at the study area in 2011. Columns represent the mean number of individuals in each age class, where error bars represent the 95% confidence interval. The vertical red line indicates the year 1975. Note, that trees below 0.3 meters tall were not detectable (dashed vertical line in b) which limited the age structure to those established before 1995 (dashed vertical line in c).

The individual tree detection algorithm identified 38,652 white spruce trees at our study area. The average observed height was 2.46 ± 1.47 m (mean ± SD) and the maximum height was 15.28 m. The stem density at the study plots was 427.07 ± 46.04 stems ha^-1^ and ranged between 358.1 – 501.34 stems ha^-1^. At the landscape level (i.e., across the ALS tiles), the average stem density was 513.12 ± 252.86 stems ha^-1^ and ranged between 100 – 1400 stems ha^-1^. The height structure at the study area revealed an exponential relationship in stems 1.4 m – 15.28 m tall (Figure 4b). However, the number of stems in each 0.1 m bin between 0.3 m (lowest detectable height) and 1.4 m tall were approximately equal.

The age structure at our study area was unimodal with a peak in the number of trees which were 36 years old in 2011 (year of aerial lidar scanning; see Section *‘Aerial lidar observations of tree height across forest stands’*) (Figure 4c), which corresponds to a tree established circa 1975 (Figure 4a). The predictions of individual tree age ranged from 16 – 182 years old. The average and median age of trees was 48.4 and 47 years old, respectively. Data could not be derived for trees less than 16 years old as these trees were < 0.3 m tall and therefore undetectable by the individual tree detection algorithm. According to age structure analyses (see e.g., Vlam et al. 2017), unimodal age structure such as that observed in Figure 4c is indicative of a population which has recently failed to recruit new individuals, so regeneration has decreased.

### 4.3 Growth rate of saplings and mature trees

The models of growth rate in saplings and mature trees were both statistically different from the null model (ANOVA, chi-squared *p* <<0.0001), confirming that growth rate did vary with age. In saplings, growth rate increased with age, but this relationship did not differ after 1975 (Table 2, Figure 5a). However, in mature trees, the relationship between growth rate and age did change significantly between the two periods (Table 2, Figure 5b). Before 1975, growth rate decreased modestly with every additional year in age (*B*_1_ = −0.2 cm^2^ year^-1^; Table 2, Figure 5b), but after 1975, growth rate increased significantly with age by 1.67 cm^2^ year^-1^ (z-test, p <0.0001). The RMSE of the sapling and mature tree models are 1.20 and 0.39 cm^2^ year^-1^, respectively.

**Figure 5.**
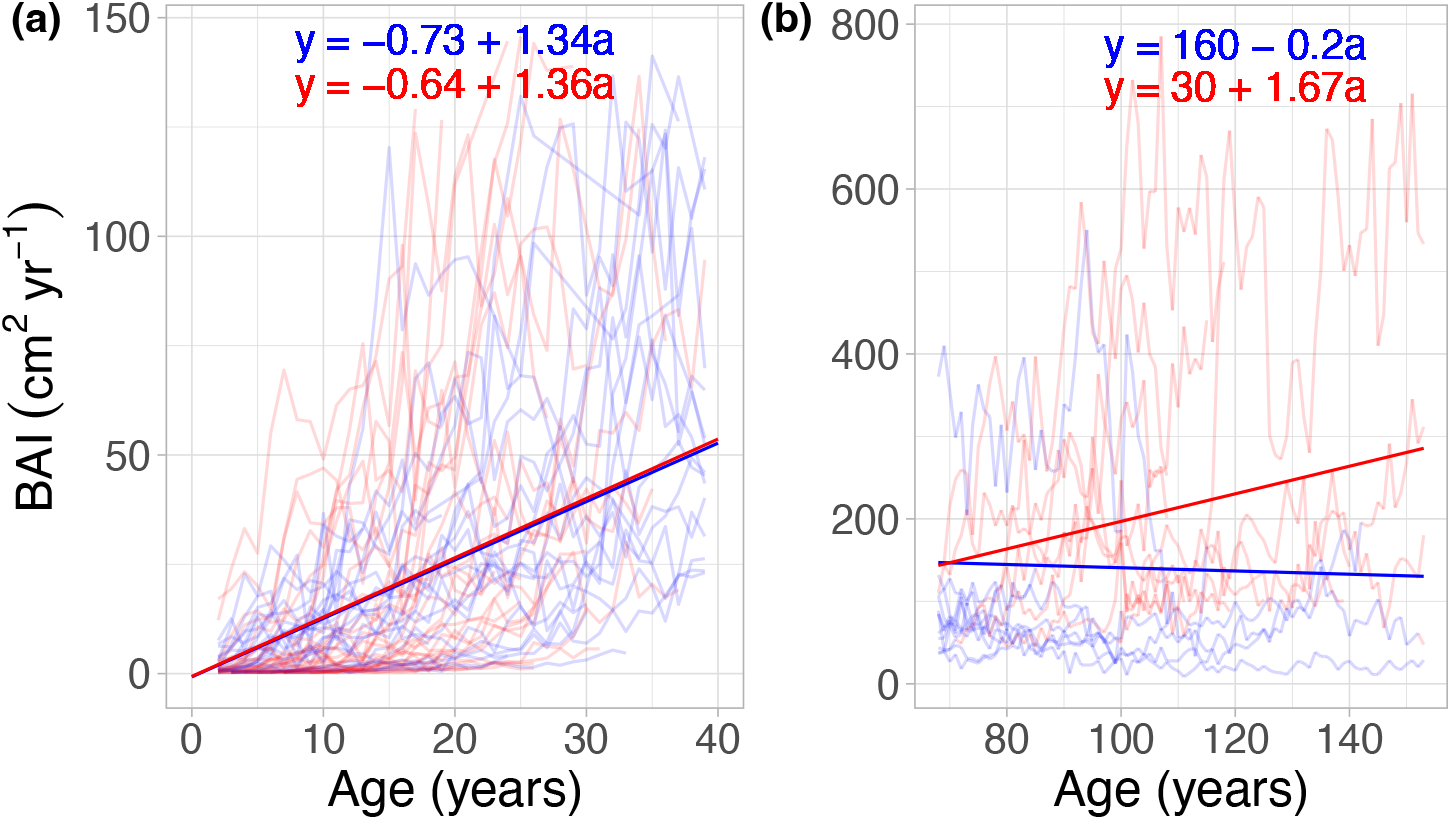
The relationship between growth rate (basal area increment, BAI; cm^2^ yr^1^) and age (years) of white spruce saplings (a) and mature trees (b) before (blue) and after (red) 1975 at Arctic treeline in the central Brooks Range, AK. The equations of the lines (i.e., Equations 2 and 3) are presented on each panel with the colors also corresponding to period. To simplify the figure and highlight the change in slope, the lines/equations present the means of the coefficients (see Table 1) and exclude the error and random effects, if applicable. The difference in slope between the two periods was only significant in mature trees (p< 0.0001; z-test).

**Table 2.** Coefficient estimates, standard error, and significance for fixed effects in the models for growth rate of white spruce saplings and trees at Arctic treeline in the central Brooks Range, AK. Bold terms are significant (p < 0.0001).

| Model | Maturity | Fixed Effect | Random Effect | Symbol | Coefficient Estimate | Std. Error | t-value | P-value<sup>1</sup> |
| --- | --- | --- | --- | --- | --- | --- | --- | --- |
| Model 1 | Saplings | Intercept | - | B<sub>0</sub> | -0.73 | 0.46 | -1.60 | 0.111 |
|  |  | **Age** |  | **B<sub>1</sub>** | **1.34** | **0.07** | **18.9** | **< 0.0001** |
|  |  | Period |  | B<sub>2</sub> | 0.09 | 0.69 | 0.13 | 0.898 |
|  |  | Age*Period |  | B<sub>3</sub> | 0.02 | 0.10 | 0.20 | 0.842 |
| Model 2 | Mature Trees | **Intercept** | **Tree ID** | **B<sub>0</sub>** | **160.4** | **22.3** | **7.20** | **< 0.0001** |
|  |  | **Age** |  | **B<sub>1</sub>** | **-0.196** | **0.035** | **-5.65** | **< 0.0001** |
|  |  | **Period** |  | **B<sub>2</sub>** | **-130.6** | **17.6** | **-7.44** | **< 0.0001** |
|  |  | **Age\*Period** |  | **B<sub>3</sub>** | **1.87** | **0.162** | **11.56** | **< 0.0001** |

<sup>1</sup> p-values derived from a t-test for saplings and from a Z-test for mature trees

### 4.4 Tree-ring width and radial stem growth vs. climate

#### Tree ring-climate analyses

The MNE tree ring chronology echoed the results of radial stem growth in mature trees: After 1975, tree ring width was 54% larger relative to previous decades (Figure 6; p < 0.0001, Wilcoxon test). The trend in radial stem growth is less prominent (7% increase; p = 0.64, Wilcoxon test) in the spline chronology due to the intrinsic nature of the detrending method. The correlations between the MNE tree-ring chronology and the monthly climate variables highlighted some prominent trends between annual growth (proxy as TRC) and environmental conditions (Table 3). Generally, warmer springs (May) and summers (June, July, August) of the current and previous years were associated with increased annual growth. Additionally, annual growth increased when the spring (May) and early- to mid-summer (June, July) of the current and previous year were drier (higher VPD) and when the previous February was wetter (lower VPD). Precipitation in July of the previous year was positively correlated with annual growth the following year. As SNO decreased in the spring (March, May) and fall (September) of the previous year, annual growth increased. Lastly, the correlation between climate index and TRC was negative (r = −0.43 ± 4.6×10^-7^, mean ± SEM; p-value < 0.0001, df = 78), suggesting that annual growth increased as the climate became warmer and drier.

**Figure 6.**
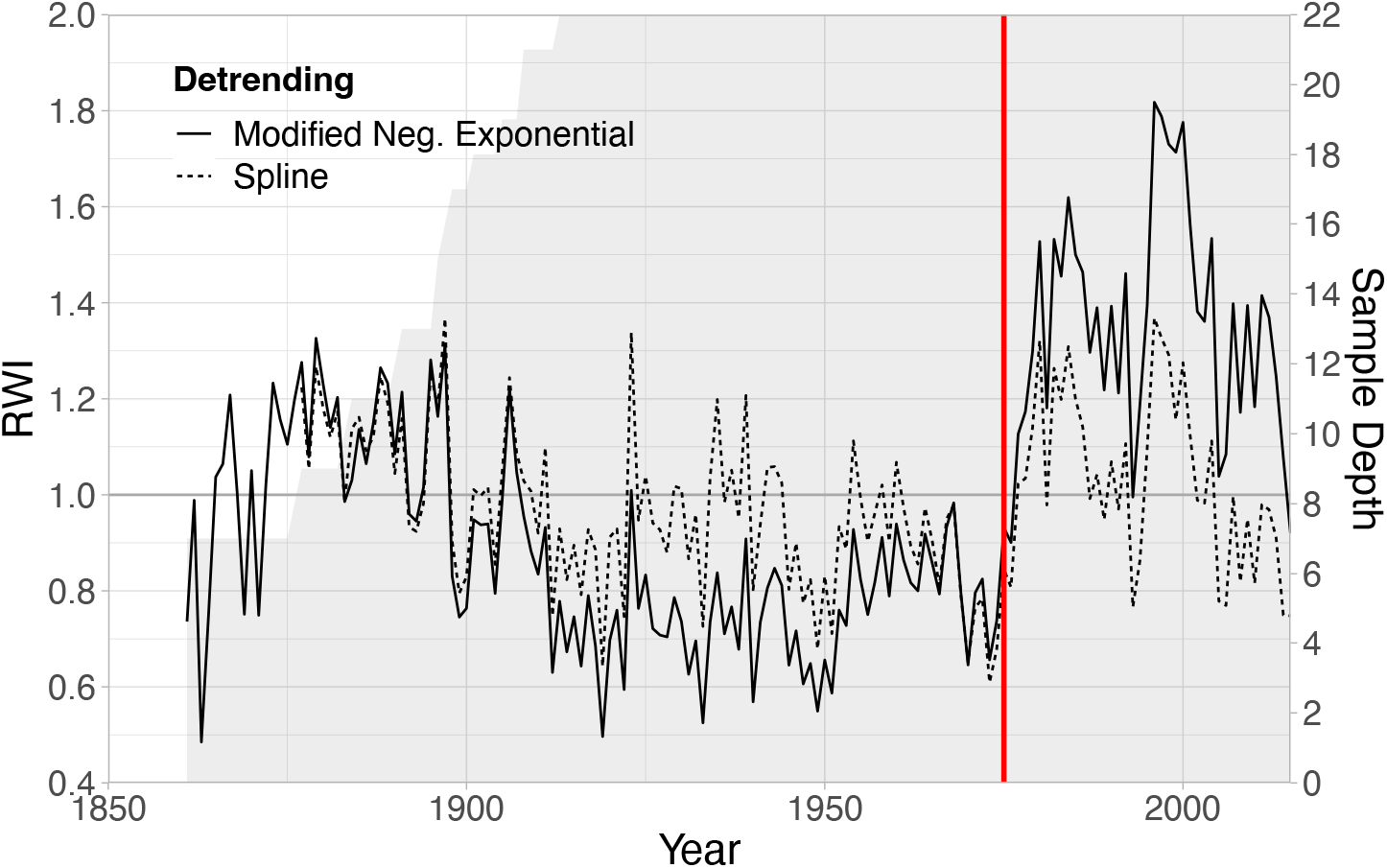
Tree-ring chronologies (TRC) with two types of detrending (modified negative exponential and two-thirds spline) of white spruce trees at Arctic treeline in the central Brooks Range, AK. Y-axis shows the ring-width index (RWI), or standardized ring width after detrending. Grey shading and secondary y-axis indicate the sample depth, or the number of cores in the chronology. The vertical red line indicates the year 1975.

**Table 3.**
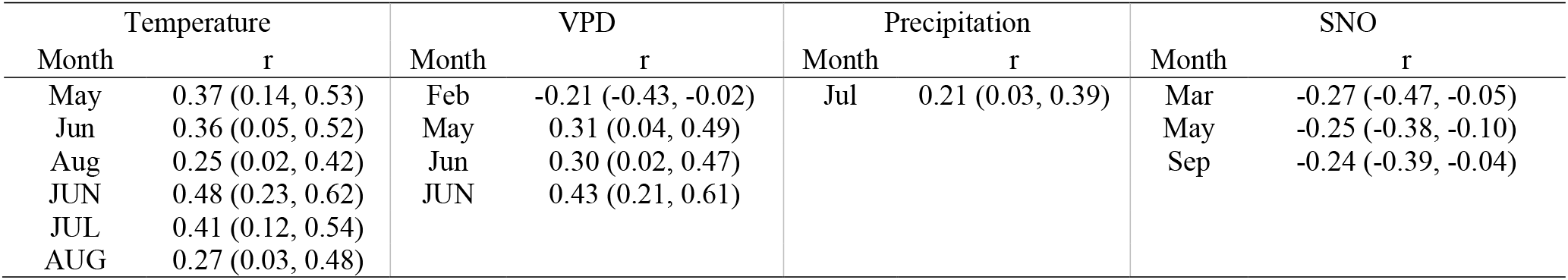
Significant Pearson correlation coefficients, r, between monthly climate and the mean negative exponential tree ring chronology (TRC) from 1900-2015 (*p* ≤ 0.01) of white spruce trees at Arctic treeline in the central Brooks Range, AK. Months of the previous year are lower case abbreviations, and the current year are in all caps. Values within parentheses indicate the 99% confidence interval around the mean.

#### Temperature divergence

The 31-year running correlations between TRC, current June temperature, and July precipitation revealed some signs of temperature divergence in our tree ring chronology. We examined the correlations with June temperature and July precipitation specifically because their correlations with the spline chronology from 1900-2015 was statistically significant (see Methods). Overall, the correlation between temperature and TRC weakened over the 20th century but was variable (Figure 7). For example, c. 1955, the correlation between TRC and current June temperature decreased and became statistically insignificant after circa 1955. After circa 1965, June temperature correlations began increasing again, only to decrease circa 1988. Meanwhile, the correlation between TRC and current July precipitation increased steadily until circa 1980 before decreasing again. The running correlations between TRC and current July precipitation were never statistically significant.

**Figure 7.**
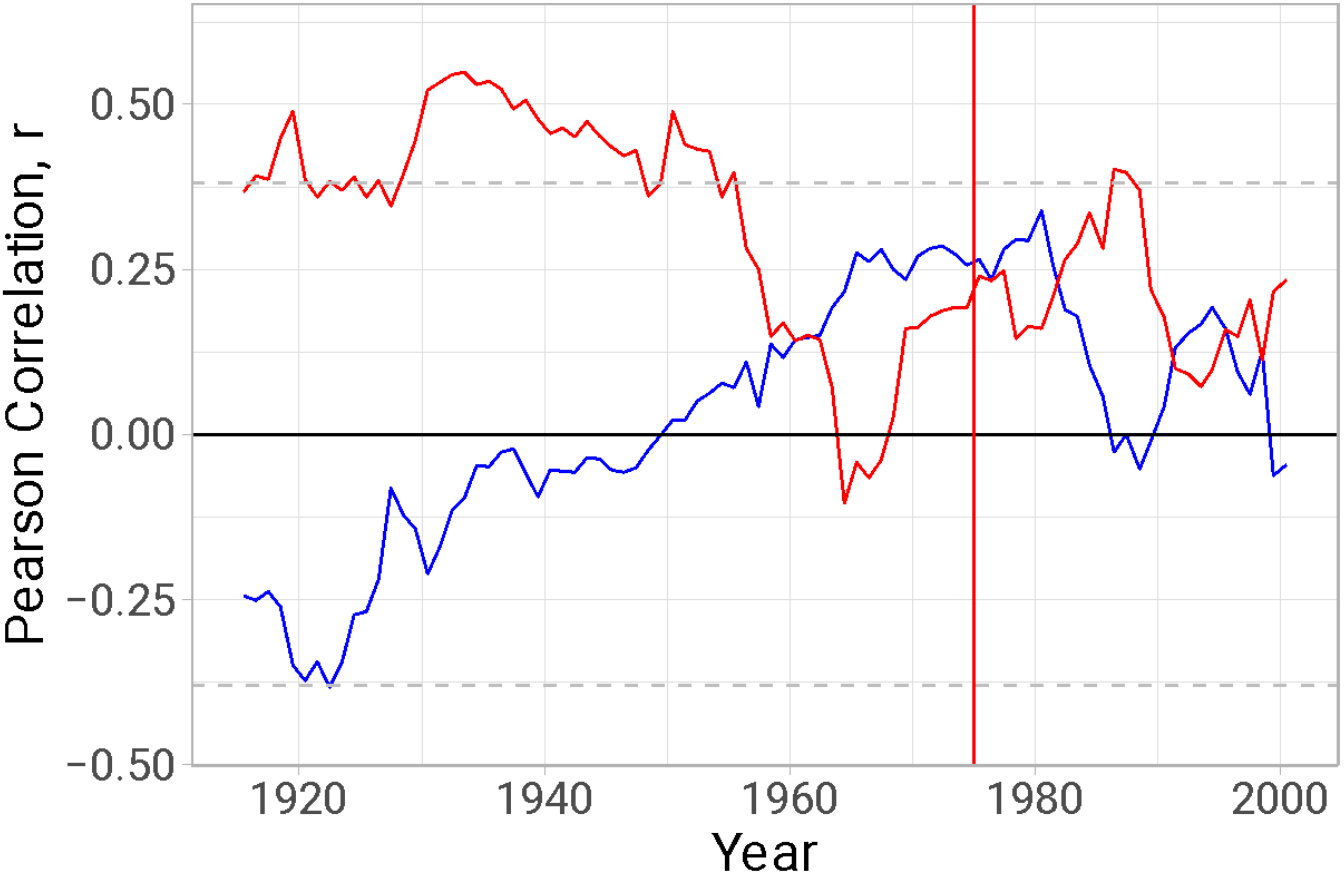
Running 31-year Pearson correlation, r, between spline-detrended TRC and mean June temperature (red line) and July precipitation (blue line) demonstrating temperature divergence of white spruce trees at Arctic treeline in the central Brooks Range, AK. The dashed, grey lines represent the 95% confidence limits, outside of which the correlation is significant. The vertical red line highlights the year 1975. Mean June temperature and July precipitation is presented because it was found to be optimal for the tree ring series analysis from 1900-2015 (see Table 3).

## 5. Discussion

Our findings reveal that climate at our latitudinal treeline site in Alaska became warmer and drier after circa 1975. Concurrently, the unimodal age structure suggests that forest regeneration decreased circa 1975 relative to previous decades, corroborating our hypothesis that warmer but drier conditions would decrease regeneration. However, in contrast to our hypothesis, our analysis of growth rate before and after 1975 showed that the warmer and drier climate did not appear to affect growth rates in saplings but increased the growth of mature trees substantially. Radial stem growth of mature trees was substantially larger after 1975 despite the overall drying trend, suggesting that mature tree growth was not or was minimally water-limited during the 20th century. However, the tree-ring chronology of mature trees exhibited positive correlations with July precipitation as well as temperature divergence starting in mid-1900s, suggesting that a shift towards water-limitation could occur if drying trends continue (D’Arrigo et al. 2008, Andreu-Hayles et al. 2011). Together, our results show divergent responses of regeneration and radial stem growth of white spruce growing at Arctic treeline in response to the same climatic stimuli.

The decrease in regeneration beginning circa 1975 was likely driven by changes in the climate. Success in early life stages relies on adequate water availability during germination (Angell and Kielland 2009), establishment (Walker et al. 2012), and seedling development (Angell and Kielland 2009) because smaller root systems make saplings highly vulnerable to water stress (Smith et al. 2003). Additionally, saplings are more likely to survive when well-covered by snow during winter since the snow pack decouples them from the harsh conditions of the free atmosphere (Körner 2012, 2016, Renard et al. 2016). While we do not have data available for tracking snow depth during the 20th century, we observed significant decreases in precipitation received as snow (SNO) during spring (i.e., −38% decrease in March and −100% decrease May after 1975) and fall (i.e., −28% decrease in September) in the years after 1975. Such a shift could have decreased the snow depth and therefore exposed seedlings to the free atmosphere earlier, ultimately decreasing survival.

While approximately half of the studied Arctic treeline sites have advanced during periods of warming (Rees et al. 2020), advance seems unlikely at our study site. The observed decrease in population regeneration at this site suggests that the tree population has not grown, and therefore treeline has not advanced. Other stationary treeline sites around the circumarctic had similar responses to climate trends such as warmer springs and summers and drier springs and falls (Rees et al. 2020). Further, the decreasing moisture gradient from western to eastern Alaska (Wilmking and Juday 2005) may explain why treeline has advanced in the western Brooks Range (Dial et al. 2022) and Seward Penninsula (Suarez et al. 1999, Lloyd and Fastie 2002) but not at our study site in the central Brooks Range. Of course, the decreased regeneration could also be explained by non-climatic factors. Many seedlings, saplings, and mature trees within the vicinity of and at our study site exhibited mild to severe browsing damage from snowshoe hares (*personal observation*). In multiple cases, snowshoe hare browsing girdled the stem and killed the tree, highlighting the strong control herbivores can have on tree survival and treeline advancement (e.g., see Olnes et al. 2018). Other factors could have affected regeneration but were beyond the scope of this study such as nutrient-availability (Sullivan 2016, Gustafson et al. 2021, Dial et al. 2022), permafrost thaw (Lloyd 2005, Maher et al. 2021), and seed dispersal (Kambo and Danby 2018a, 2018b, Dial et al. 2022). While we cannot conclude that climate was the main driver in decreasing regeneration after 1975, the striking synchronicity of the shift in climatic conditions and the shift in regeneration - plus the knowledge demonstrated from prior research that climate constrains regeneration in tree populations - strongly suggests a mechanistic link.

In contrast to the decrease in regeneration of saplings, the growth rates of mature trees increased in response to warming and drying after circa 1975. This finding contradicts our hypothesis that both growth and regeneration would decrease in response to warming and drying. However, these differing responses may be explained by studies which show that climate change influences saplings and mature trees differently, often because of their difference in size (Körner 2012, 2016, Stovall et al. 2019). For example, larger, mature trees may be able to grow under drier conditions than saplings because of their more developed root systems which can more easily access deeper water reserves (Körner 2012, Kambo and Danby 2018a, 2018b). While it appears that the growth response of mature trees at our site is not currently water-limited, the presence of temperature divergence in our tree ring chronology suggests that continued drying and warming may cause future water-limitation to impede growth (Peters et al. 2021).

Surprisingly, our results show that relative to before 1975, the growth of saplings did not change significantly after 1975. At face value, one might conclude that the growth of saplings was not impacted by warming and drying conditions, although seedling and sapling mortality were not measured and may confound this conclusion. This presents a question for future research: How and why would warming and drying conditions affect regeneration but not sapling growth? Recent studies suggest that sapling survival depends on overcoming higher respiratory carbon losses (Griffin et al. 2021) which may in turn affect growth. Furthermore, the carbon balance constraints resulting from high respiratory rates and low photosynthetic rates are exaggerated at treeline relative to the southern range limit of white spruce (Griffin et al. 2022, Schmiege et al. 2022). Future research is necessary to identify the physiological explanations for this finding and might consider investigating the growth and carbon balance of both small and large spruce trees growing in different water regimes.

The age-height model enabled us to predict the age of trees detected by aerial-lidar and therefore expand our scope to the landscape level covering 250 ha. The study of such large territory along the treeline was only possible due to the implementation of the new methodological approaches presented here. Tree height is not typically an accurate proxy for tree age due to several factors which can stunt or accelerate the growth of individuals such as gap dynamics or environmental stress (Kuuluvainen et al. 2002). However, in this population we validated the relationship and found a strong relationship between age and height (r^2^ = 0.82, RMSE = 23.8 years). More research is required to understand this unusually strong relationship in trees; however, it is likely because trees at our study site are well spaced, therefore the effects of gap dynamics and competition are limited. We do observe an increase in the variation of age among mature trees, possibly due to variation in environmental stress. When studying other species or older populations, we recommend verifying age-height models, as the mature trees in this study are still relatively young in a general context.

Future research may advance the method presented here by improving the predictive accuracy of the age-height model, diversifying the species in the model, and using lidar with improved vertical accuracy to decrease the height detection threshold of individual trees. Adding other predictor variables to the age-height model, such as diameter at breast height or distance to nearest neighbor, could greatly improve model accuracy. Additionally, developing age-height models for other treeline species would make our methods more widely applicable. Lastly, using higher resolution aerial lidar scans (> 8 returns per m^2^) and higher vertical accuracy in future studies would enable the observation of trees below the height detection threshold (i.e., 30 cm) and decrease the possibility that small shrubs (i.e., > 30 cm tall) were misclassified as spruce seedlings and vice versa. If small trees were misclassified as shrubs, then regeneration could be higher than reported. Conversely, if shrubs were misclassified as small trees in the sample population assessed in this study, regeneration was even more negatively affected by climate.

### Conclusion

The results of our study show that at our Arctic treeline site in interior Alaska, shifts towards a drier and warmer climate were highly associated with a decrease in regeneration of saplings and increased growth of mature trees. Together, this lends evidence that this section of treeline has remained stationary since circa 1975 while mature tree growth has increased. However, only one or two successful cohorts are necessary to advance treeline position (Körner 2016). Should the many stochastic factors necessary for increased regeneration temporarily align, this section of the Arctic treeline may advance. Therefore, we stress that continued monitoring is necessary to properly assess current treeline behavior. Our novel method for monitoring regeneration offers a means to do so that is less labor-intensive than traditional methods. Wide application of this method has the potential to open research to less well-documented locations of the Arctic treeline and clarify the varied responses of Arctic treeline to climate change.

## 6. Acknowledgements and Author Contributions

Funding for this research came from NASA Terrestrial Ecology grant NNX15AT86A, NASA ABoVE grant (Eitel-01), and NSF grant OPP21-24885. JEJ, KLG, NB, JUHE, and LAV developed the initial framework and objectives of the study. JEJ, KLG, NB, JUHE, LAV, and AJM assisted in data collection in the field. CS extracted tree location and height using the ITD algorithm. Tree ring analysis was led by JEJ and RO. RD and LAH were partly supported by NSF OISE-1743738 and NSF-2124885. JEJ drafted the initial version of the manuscript. All coauthors contributed to the interpretation of the results and revisions of the manuscript. We thank Hélène Genet at the University of Alaska Fairbanks for providing us with historical climate data of our study site via the Scenarios Network for Alaska and Arctic Planning (SNAP) database. We are also grateful for the support provided by the staff of the ABoVE Fairbanks logistics office, notably Sarah Sackett. We would also like to give a special thanks to Sarah Bruner and Jyoti Jennewien for assistance in the field; Arjan Meddens for assistance with geospatial analysis; Mukund Rao for advice in analyzing tree rings; and Duncan Menge for statistical advice and collaboration.

## 7. Data Availability Statement

The data for this study has been submitted to Oak Ridge National Laboratory (ORNL) Distributed Active Archive Center (DAAC) and is in the final stages of reviewing. Data will be published prior to the publication of this study.

