## Supplementary Material Table 1 for "Growth increases but regeneration declines in response to warming and drying at Arctic treeline in white spruce (*Picea glauca*)"

Supplementary Material Table 1. Variance of Random Effects in Growth Rate v.s. tree age model in mature trees before and after 1975.

| Groups | Name | Variance | Std. Dev. |
|---|---|---|---|
| Tree ID | Intercept | 2273.7 | 47.7 |
| Residual | | 0.16 | 0.40 |
